# Connexin 43 K63-polyubiquitination on lysines 264 and 303 regulates gap junction internalization

**DOI:** 10.1101/211607

**Authors:** Rachael M. Kells-Andrews, Rachel A. Margraf, Charles G. Fisher, Matthias M. Falk

## Abstract

Gap junctions (GJs) assembled from connexin (Cx) proteins play a pivotal role in cell-to-cell communication by forming channels that connect the cytosols of adjacent cells. Connexin 43, the best-studied Cx, is ubiquitously expressed in vertebrates. While phosphorylation is known to regulate multiple aspects of GJ function, much less is known about the role ubiquitination plays in these processes. Here we show by using ubiquitination-type specific antibodies and Cx43 lysine (K) to arginine (R) mutants that a portion of Cx43 in GJs can become K63-polyubiquitinated on K264 and K303. Relevant Cx43 K/R mutants assembled significantly larger GJ plaques, exhibited much longer protein half-lives and were internalization impaired. Interestingly, ubiquitin-deficient Cx43 mutants accumulated as hyper-phosphorylated polypeptides in the plasma membrane, suggesting that K63-polyubiquitination may be triggered by phosphorylation. Phospho-specific Cx43 antibodies revealed that upregulated phosphorylation affected serines 368, 279/282, and 255, well-known regulatory PKC and MAPK phosphorylation sites. Together, these novel findings suggest that upon internalization, some Cx43 in GJs becomes K63-polyubiquitinated, ubiquitination is critical for GJ internalization, and that K63-polyubiquitination may be induced by Cx phosphorylation.

**Summary Statement:** Here we show that connexin 43 in gap junctions becomes K63-poly ubiquitinated on lysines 264 and 303 and its requirement for gap junction endocytosis. These novel findings significantly contribute to our understanding of GJ turnover and patho-/physiology.

**Abbreviations used:**

AGJ: annular gap junction
AMSH: associated molecule with the SH3 domain of STAM
CME: clathrin-mediated endocytosis
Cx: Connexin
Cx43: Connexin 43
DUB: deubiquitinase
GJ: gap junction
MonoUb: monoubiquitin
Nedd4-1: neural precursor cell expressed developmentally down-regulated protein 4-1
PM: plasma membrane
PolyUb: polyubiquitin
TPA: 12-O-Tetradecanoylphorbol 13-Acetate
TX-100: Triton X-100
RT: room temperature
Ub: ubiquitin

## INTRODUCTION

Ubiquitination, the addition of a small, 76 amino acid, 8.5 kDa protein to a target protein is known to play an important function in protein degradation (Komander and Rape, 2012; Nguyen et al., 2013; Ravid and Hochstrasser, 2008). Multiple types of ubiquitination (mono- and several different types of poly-ubiquitination) are known to execute these functions (Komander and Rape, 2012). Ubiquitin (Ub) is covalently attached to lysines (K) on target proteins by an enzyme cascade consisting of an E1 (Ub-activating), E2 (Ub-conjugating), and E3 (Ub-ligating) enzyme (Chen and Sun, 2009). Ub can also be removed from its target via proteases termed de-ubiquitinases (DUBs). Ub has seven internal lysines (K6, K11, K27, K29, K33, K48 and K63), all of which (and also the N-terminal methionine) are capable of forming linkages to the C-terminal glycine of subsequent Ub moieties, forming polyubiquitin (polyUb) chains. The two best studied polyUb chains are K48-polyUb, which is known to lead to proteasomal degradation, and K63-polyUb, which-besides other functions- is known to signal endo- and phago-lysosomal degradation (Komander and Rape, 2012). Ubiquitination of gap junction (GJ) proteins, termed connexins (Cxs), has also been reported (Girao et al., 2009; Laing and Beyer, 1995; Laing et al., 1997; Leithe and Rivedal, 2004; Ribeiro-Rodrigues et al., 2014), however published results are inconsistent suggesting that diverse types of ubiquitination with different functions may occur at individual stages of the connexin life-cycle.

Connexins (Cxs) are the four-pass transmembrane protein components of GJs that serve as a pathway for intercellular communication by physically coupling cells and allowing the passage of small metabolites, signaling molecules, and ions. Cxs have two extracellular loops, one intracellular loop, and an intracellular N- and C-terminus. Six Cxs oligomerize to form a hemichannel or connexon, which is trafficked to the plasma membrane (PM) and docks with a hemichannel of an adjacent cell forming the complete GJ channel. Accrual of multiple channels at the PM into clusters or two-dimensional arrays forms typical GJ plaques. Interestingly, once hemichannels dock at the PM, they can no longer be physiologically separated (Goodenough and Gilula, 1974). Therefore, in a process that requires the clathrin-mediated endocytic (CME) machinery, one of the two adjacent cells invaginates the GJ forming an annular gap junction (AGJ vesicle or connexosome) in the cytoplasm of one of the coupled cells (Falk et al., 2009; Gaietta et al., 2002; Jordan et al., 2001; Lauf et al., 2002; Piehl et al., 2007). Directionality may be achieved by phosphorylation-induced removal of a scaffolding protein, ZO-1, from the GJ plaque surface in the acceptor cell (Gilleron et al., 2009; Thevenin et al., 2017). The AGJ is then trafficked for degradation by autophago-lysosomal (under physiological, pathological, and starvation conditions) (Bejarano et al., 2012; Fong et al., 2012; Hesketh et al., 2010; Lichtenstein et al., 2010) and possibly endo-lysosomal pathways under TPA (12-O-Tetradecanoylphorbol 13-Acetate, a Diacylglycerol analog) treatment (Fykerud et al., 2012; Leithe et al., 2009). In humans, there are 21 different Cx proteins, which are identified by molecular weight. Cx43 is the best-studied Cx and is expressed in most tissues. Mutations in Cx43 have been linked to heart failure, ischemia, and hypertrophy (Fontes et al., 2012), and the developmental bone malformation, Oculodentodigital Dysplasia (ODDD) (Batra et al., 2012). Proper regulation of Cx43 trafficking and turnover is therefore imperative for human health and disease.

Cx43 ubiquitination was initially discovered by Laing and Beyer in 1995 and characterized as a signal leading to Cx43 degradation by the proteasome (Laing and Beyer, 1995). Later, the same group published evidence suggesting that ubiquitination plays a role in both, proteasomal and lysosomal degradation (Laing et al., 1997). Afterward, it was reported that Cx43 likely is modified by multiple mono-ubiquitinations (Girao et al., 2009; Leithe and Rivedal, 2004), while more recent evidence suggests that Cx43 interacts with AMSH (associated molecule with the SH3 domain of STAM) (Ribeiro-Rodrigues et al., 2014). AMSH is a DUB with specificity toward K63-polyubiquitinated proteins (McCullough et al., 2004; Sato et al., 2008), suggesting that Cx43 may also become K63-polyubiquitinated (Ribeiro-Rodrigues et al., 2014). Likely, the different discovered types of Cx43 ubiquitination distinguish the degradation of single, misfolded Cx polypeptides (by the 26S proteasome) from the degradation of GJs and AGJs (by phago- and endo-lysosomal pathways). However a rigorous analysis of where and when ubiquitination occurs on Cxs and at which stages of its ‘life-cycle’ (Cx polypeptides, connexons, GJs, AGJs) has not been performed; nor is it known what signals trigger potential Cx43 ubiquitination and whether Cx43 ubiquitination may have additional functions beyond its proteasomal degradation.

Here, we used Ub type-specific antibodies and generated Cx43 lysine to arginine mutations to investigate whether Cx43 ubiquitination occurs in GJs, what type of ubiquitination may occur, and what its role might be for GJ function. We found that apparently two juxtaposed lysine residues (K264, K303) in the C-terminal domain of Cx43 become K63-polyubiquitinated, and that ubiquitination is required for constitutive GJ internalization. Interestingly, K63-polyUb-deficient Cx43 accumulated as hyper-phosphorylated polypeptides in the PM, suggesting that phosphorylation may trigger Cx43 ubiquitination.

## RESULTS

### K63- but not K48-polyubiquitin specific antibodies colocalize with plasma membrane and internalized gap junctions

To determine whether and what type of ubiquitination may occur on Cx43 in GJs, we immunostained endogenously expressed Cx43 in primary porcine pulmonary artery endothelial cells (pPAECs) with antibodies directed against Cx43 and different types of ubiquitin (**Figure 1**). FK2, which recognizes both monoUb and polyUb, colocalized with Cx43 at GJ plaques (insert 1, marked with arrows) and internalized, annular GJs (AGJs) (insert 2, marked with arrowheads) (**Figure 1A**). Next, we co-immunostained Cx43 with antibodies that detected only polyubiquitinated proteins (FK1). We found that colocalization of Cx43 GJ plaques and AGJs with FK1 was also evident (**Figure 1B**). To test whether the detected polyubiquitin modification includes K63-based polyUb, we conducted co-immunostaining with an antibody that detects only K63-polyUb (HWA4C4). Again, robust yet more restricted colocalization of GJs and AGJs with the K63-polyUb antibody was detected (**Figure 1C**). To investigate whether K48-polyubiqitination (a known proteasomal degradation signal) may occur in GJs, we co-immunostained PAECs with an antibody that detects only K48-polyUb. No evidence of colocalization with K48-polyUb was found on Cx43 GJs and AGJs, suggesting that connexins and oCx-binding proteins in GJs are not K48-polyubiquitinated (**Figure 1D**). Interestingly, all three Ub-specific antibodies (FK1, FK2, and HWA4C4) localized to defined GJ plaque areas, suggesting that active, endocytic activity may occur specifically in these areas; a hypothesis that is further supported by the colocalization of these ubiquitin antibodies also with internalized AGJ vesicles. Quantitation of ubiquitin colocalizing with GJs in the PM varied from app. 60 to 10 *%* for FK2, FK1, and HWA4C4 depending on stringency of threshold settings, while no colocalization was observed with Apu2 under any of these conditions. Taken together, our results indicate that Cx43 in GJs, or connexin-associated proteins in GJs and AGJs can become K63-polyubiquitinated. Our finding correlates with the enzyme specificity of the E3 ubiquitin ligase, Nedd4-1 (neural precursor cell expressed developmentally down-regulated protein 4-1), which preferentially K63-polyubiquitinates its substrates (Kim et al., 2007), and has been found to ubiquitinate Cx43 (Girao et al., 2009; Leykauf et al., 2006). Our data also correlates with recent biochemical evidence suggesting that Cx43 interacts with the K63-polyUb specific DUB, AMSH (Ribeiro-Rodrigues et al., 2014).

**Figure 1:**
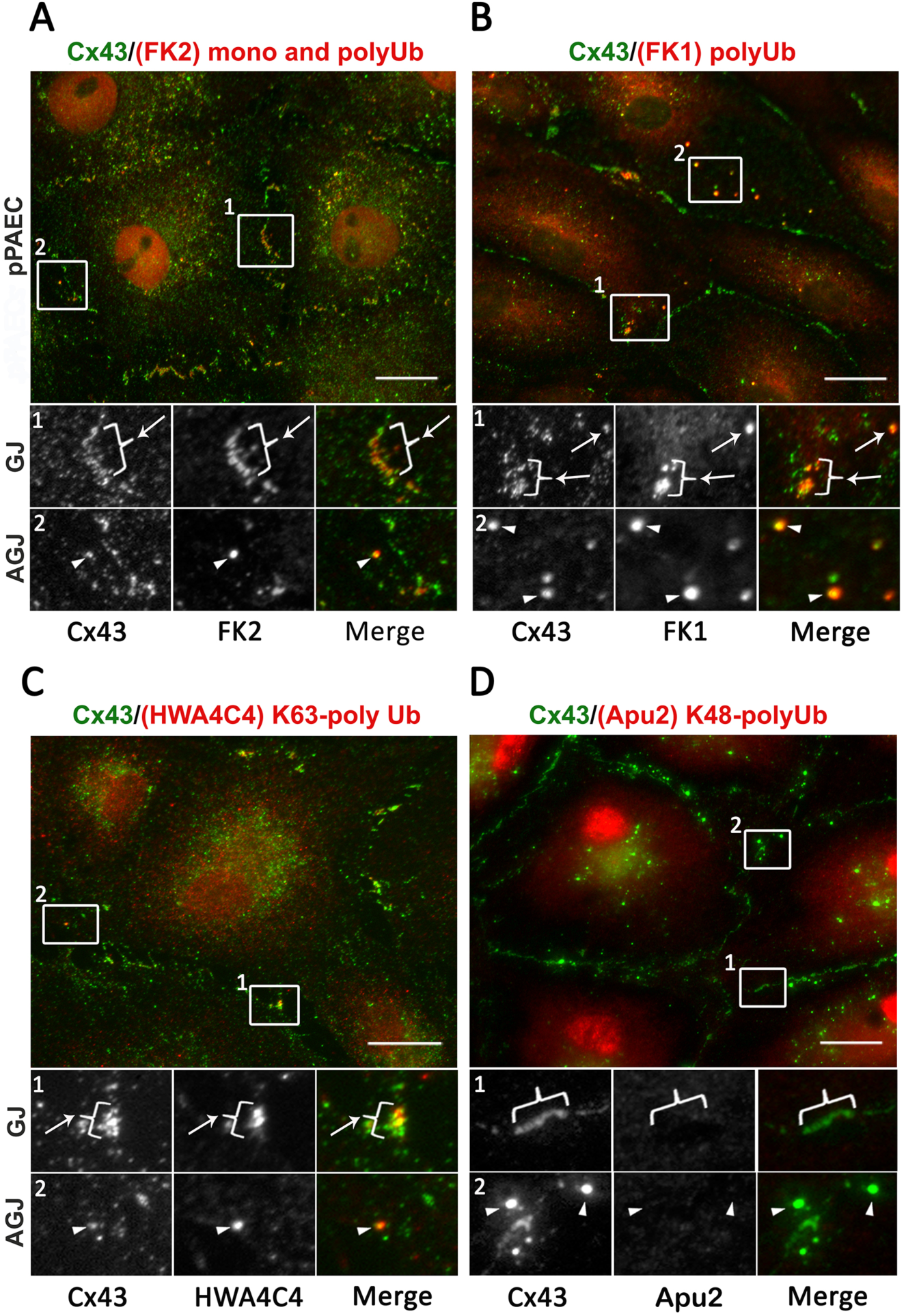
Ubiquitin-specific antibodies colocalize with Cx43 in GJs and AGJs. Endogenously Cx43-expressing primary porcine pulmonary artery endothelial cells (pPAECs) were immunostained with antibodies directed against Cx43 (green) and ubiquitin specific antibodies (red). Below each panel, magnified insets highlight GJs (brackets and arrows) and AGJs (arrowheads). Cx43 antibodies robustly colocalized with mono and polyUb specific antibodies (FK2) **(A)**, only polyUb specific antibodies (FK1) **(B)**, and with K63-polyUb specific antibodies (HWA4C4) at distinct areas of GJs and with AGJs **(C)**, but not with K48-polyUb specific antibodies (Apu2) **(D)**, suggesting that Cx43, or a Cx43-binding protein became K63-polyubiquitinated. Images and blots shown in this and the following figures are representative of at least 3 independent experiments. Scale bar = 20 μm.

### Cx43 in GJs can become K63-polyubiquitinated

To further investigate whether K63-polyubiquitination in GJs occurs on Cx43 itself, or on a Cx43-associated protein, we performed Triton X-100 (TX-100) insolubility assays with endogenously Cx43 expressing pPAECs and exogenously Cx43 expressing MDCK cells (not expressing endogenous Cxs) (Dukes et al., 2011) to separate the TX-100 insoluble (GJs, AGJs) from the soluble (Cxs, connexons) fractions using a well-established method (Musil and Goodenough, 1991). MDCK cells were chosen in this and all other experiments as exogenous expression system (with the exception of GJ plaque size analyses, see below) as high transfection efficiencies (>75%) were achieved in this cell line. TX-100 soluble and insoluble fractions were analyzed via western blot and probed for Cx43 (**Figure 2A**). In both fractions, a well-characterized triplet of Cx43 bands was detected corresponding to the fastest migrating form of Cx43 (P0) and the slower migrating phosphorylated forms of Cx43, commonly referred to as P1 and P2. TX-100 insoluble fractions contained a number of additional, higher molecular weight bands migrating above Cx43 and appearing as a smear in some fractions (marked with brackets in A to C) that potentially corresponded to ubiquitinated Cx43 polypeptides. Two bands migrating around 70 kDa and 150kDa were especially prominent in most fractions (marked with asterisks in A to C). When TX-100 insoluble fractions of cells untreated or treated with 20 μM PR-619, a pan-DUB inhibitor, for 1.5 hours prior to lysis were probed for monoUb/polyUb (FK2), polyUb (FK1), and K63-polyUb (HWA4C4), dramatically increased levels of ubiquitinated proteins were detected in the analyzed fractions that migrated as a ladder or a smear of higher molecular weight proteins together with the bands migrating at around 70 and 150 kDa that also increased in the presence of PR-619 (**Figure 2B**). To provide further evidence whether Cx43 may become K63-polyubiquitinated, MDCK cells were transfected transiently with wild type Cx43 cDNA, lysed and fractionated as described above for pPAECs 24 hours post transfection. The TX-100 insoluble pellet was then subjected to K63-polyUb and Cx43 immunoprecipitations. Higher molecular weight bands, including the 70 and 150 kDa bands were detected in the GJ-containing pellet fraction when either Cx43 was precipitated and probed with K63-polyUb specific antibodies (**Figure 2C, left panel**), or K63-polyubiquitinated proteins were precipitated and probed with Cx43 specific antibodies (**Figure 2C, right panel**), suggesting that Cx43 itself in GJs can become K63-polyubiquitinated. That the higher molecular weight signals corresponded to ubiquitinated Cx43-binding partner(s) is not likely as protein complexes containing Cx43 should have been dissociated in the SDS and β-mercaptoethanol-containing SDS-PAGE sample buffer. Protein A-Sepharose beads alone and immunoprecipitation reactions without antibodies analyzed in parallel did not precipitate any proteins. It should be noted that the amount of detected higher molecular weight bands is small compared to the total level of non-ubiquitinated Cx43 detected in the insoluble fractions (**Figure 2A**, right panel). This is likely due to ubiquitination being a transient process that probably affects only a relatively small portion of Cx43 in a GJ plaque (the portion of the plaque that internalizes or triggers internalization). In addition, only some of the connexins in a connexon may become ubiquitinated. This is indicated by the fact that unmodified Cx43 was detected together with ubiquitinated Cx43 when total K63-ubiquitinated proteins were precipitated and probed with Cx43 specific antibodies (Figure 2C, right panel, band marked with Cx43); while unmodified Cx43 was not detected when Cx43 was precipitated and probed with K63-polyUb specific antibodies (Figure 2C, left panel). Together with our immuno-colocalization analyses shown in Figure 1 these data suggest that Cx43 in GJs and probably also in AGJs, both in endogenously as well as in exogenously Cx43 expressing cells can become K63-polyubiquitinated.

**Figure 2:**
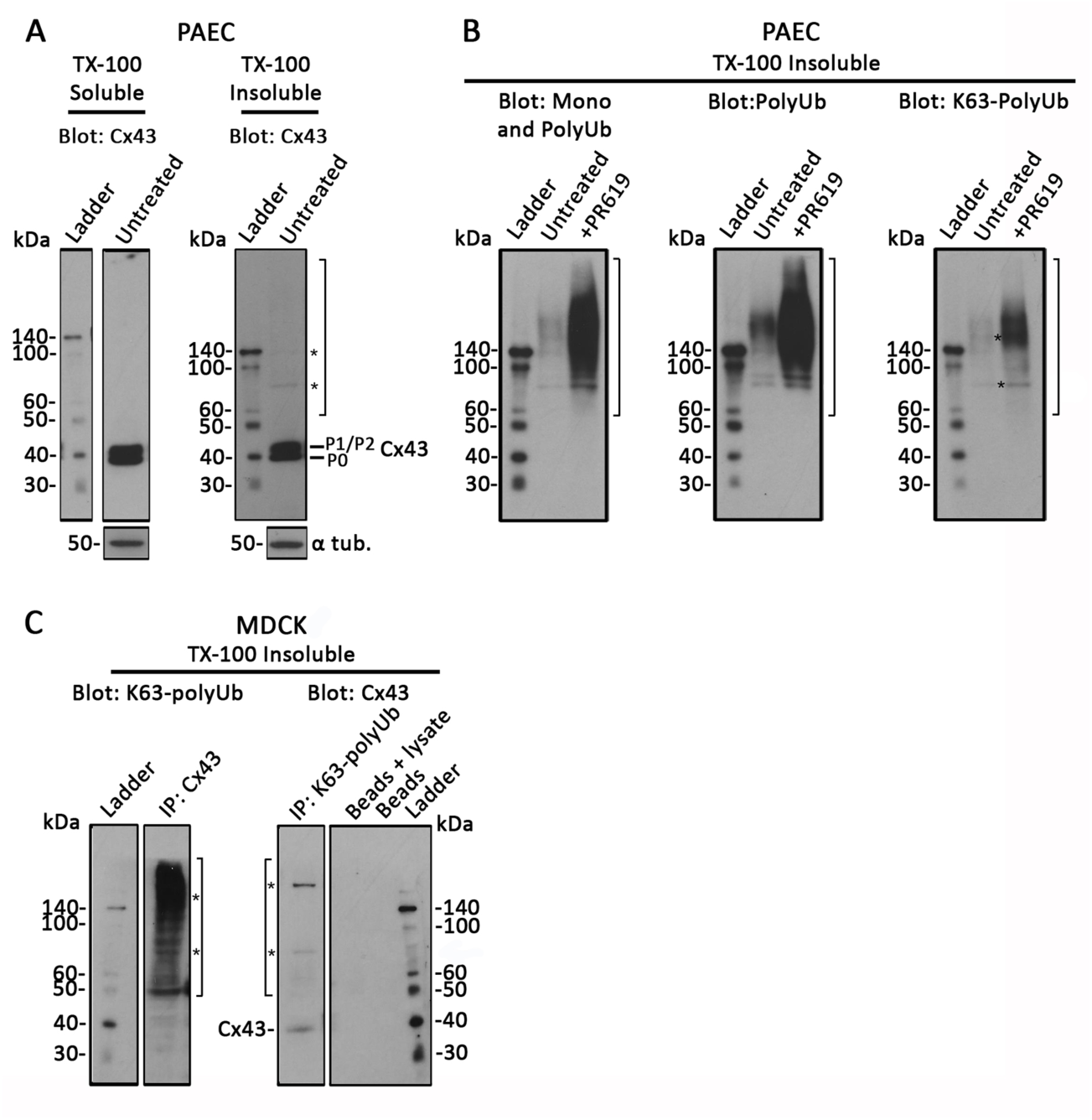
Both, Cx43- and K63-polyUb specific antibodies precipitate Cx43-higher molecular weight forms. Endogenously Cx43-expressing pPAECs (in **A** and **B**), and exogenously Cx43 expressing MDCK cells (in **C**) were lysed in buffer containing 1% TX-100, separated into TX-100 insoluble (GJ/AGJ) and soluble (Cx, connexons) fractions, probed using Cx43 and ubiquitin specific antibodies, and analyzed by western blot. **(A)** Mature forms of Cx43 (P0, P1, P2) migrating at the predicted molecular weight of app. 43 kDa were detected in both, TX-100 soluble and insoluble fractions. In addition, small amounts of higher molecular weight forms (bracket) including two species migrating at app. 70 and 150 kDa (asterisks) were detected in the insoluble fraction in this and the analyses shown in B and C, suggestive of potentially ubiquitinated Cx43. **(B)** PAECs were treated or not with the DUB inhibitor, PR-619, TX-100 extracted, and probed with mono/polyUb (FK2), polyUb (FK1), and K63-polyUb (HWA4C4) specific antibodies. In all instances, a smear of higher molecular weight bands (bracket) together with species migrating at app. 70 and 150 kDa (asterisks) were detected with all Ub-specific antibodies (and e.g. with K63-polyUB antibodies), especially in the lysates derived from cells in which protein-deubiquitination was blocked. **(C)** Cx43 and K63-polyubiquitinated proteins present in TX-100 insoluble fractions were precipitated from transfected MDCK cells and probed for K63-polyUb proteins and Cx43, respectively. Resulting blots again revealed a mixture of higher molecular weight bands (brackets) together with species migrating at app. 70 and 150 kDa (asterisks), further suggesting Cx43 K63-polyubiquitination. Note that mature Cx43 migrating around 43 kDa was also detected when K63-polyubiquitinated proteins were precipitated and probed with Cx43 specific antibodies (right panel), but not in the inverse experiment (left panel), suggesting that only a portion of Cx43 subunits in connexons are ubiquitinated.

### Cx43 GJs with critical lysine residues mutated to arginines accumulate in the plasma membrane

To determine which lysines in Cx43 GJs may become K63-polyubiquitinated, we transiently transfected HeLa cells with rat Cx43 constructs containing a set of C-terminal lysine (K) to arginine (R) mutations (called K/R) in order to block ubiquitination. We focused on mutating lysines in the C-terminus of Cx43 as this domain is highly post-translationally modified and is known to interact with numerous binding partners (reviewed in Thevenin et al., 2013). Mutant 6K/R contains six lysines within the C-terminus (K258, K264, K287, K303, K345, and K346) mutated to arginines (**Figure 3A**, boxed blue and green). Mutant 3K/R contains the central lysines K264, K287, and K303 mutated to arginines (**Figure 3A**, boxed green only). 24 hours post transfection, cells were fixed and immunostained using Cx43 antibodies. Both mutants trafficked to the plasma membrane and assembled into GJ plaques with equivalent efficiency compared to wild type, however cells expressing either 6K/R or 3K/R mutants formed noticeably larger GJ plaques (**Figure 3B, left,** shown for transfected HeLa cells). Total GJ plaque area in WT and mutants was quantified by outlining GJ plaques and recording total fluorescence intensity of the outlined regions. Average fluorescence intensity and corresponding GJ plaque size per cell pair significantly increased by approximately 1.5-2 fold in both mutants (6K/R and 3K/R increased to 8.60 ×10^6^ ± 0.65 and 10.20 × 10^6^ ± 0.85 arbitrary units, respectively; n = 3) when compared to WT (5.35 × 10^6^ ± 0.58 arbitrary units) (**Figure 3B, right**). 6K/R and 3K/R mutants gave similar results, implying that the additional peripheral residues in the 6K/R mutant (K258, K345, and K346), or any other of the lysine residues that are present in the Cx43 polypeptide (**Figure 1A**, intracellular located lysines labeled in red) did not become ubiquitinated. HeLa cells were used in this experiment, as they in general form larger, easier to quantify GJs due to the more spread-out morphology of Hela compared to MDCK cells. Moreover, a comparable result was obtained when equal amounts of cell lysates were analyzed and probed for Cx43 protein content in Cx-deficient MDCK cells expressing wild type and mutant Cx43 (**Figure 3C**). Quantification of 6K/R and 3K/R mutants resulted in 8.88 ± 2.70 and 7.71 ± 2.95 fold increases in total Cx43 polypeptide amounts compared to wild type, respectively (**Figure 3C**). Taken together, these results indicate that mutating critical lysine residues in the Cx43 C-terminal domain causes accumulation of Cx polypeptides and GJs in the PM, that based on the known role of K63-polyubiquitination suggests impairment of GJ internalization.

**Figure 3:**
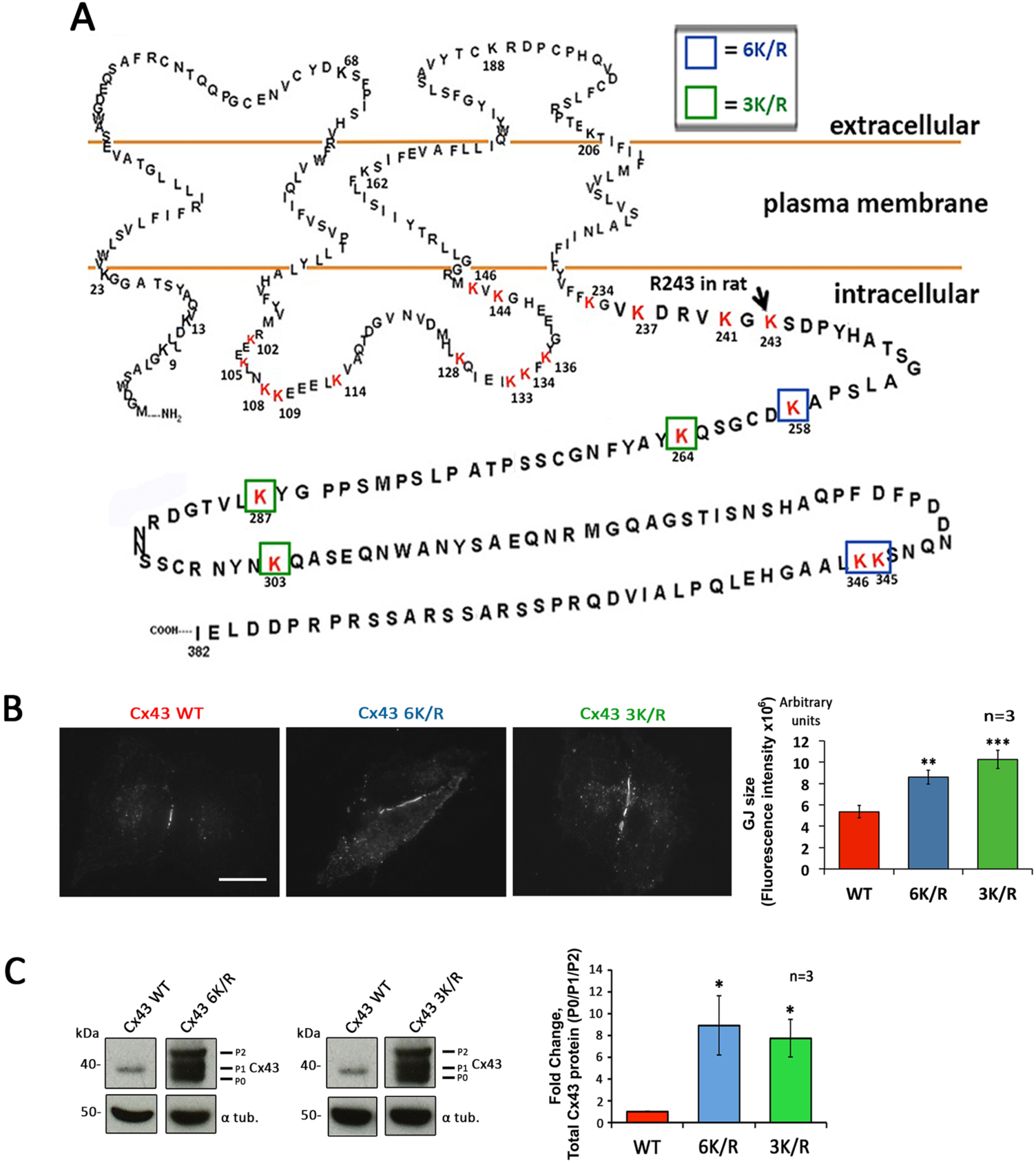
Cx43 mutants with lysine residues mutated form larger GJ plaques. (A) Amino acid sequence of human Cx43 with lysines (K) in the intracellular loop and the C-terminus potentially accessible to ubiquitination in GJs highlighted in red. Lysines mutated to arginines (R) in the C-terminus of Cx43 are boxed in blue and green for mutant 6K/R, and green only for mutant 3K/R. **(B)** Cx-deficient HeLa cells (forming larger GJ plaques compared to PAEC and MDCK cells used in all other analyses) were transfected with wild type or K/R mutants, immunostained with anti-Cx43 followed by Alexa488-conjugated secondary antibodies, and GJ length and number determined. Fluorescence intensity of outlined GJs (in 73, 98 and 83 cell pairs expressing Cx43 wild type, 6K/R, and 3K/R, respectively) representative of GJ size was measured and blotted. Both K/R mutants showed significantly larger GJ plaques compared to wild type Cx43. **(C)** Total Cx43 protein levels of wild type, 6K/R and 3K/R mutants exogenously expressed in MDCK cells were analyzed by western blot. In both mutants, app. 8 times the amount of Cx43 including a large amount of phosphorylated P1 and P2 variants was detecte. Lanes of blots are separated to indicate extraneous lanes between wild type and 6K/R, and wild type and 3K/R, however lanes were derived from the same gels. Scale bar = 20μm.

### K63-polyubiquitin-deficient Cx43 mutants exhibit significantly longer protein half-lives

To further investigate the inhibitory effect of mutating lysine residues on GJ internalization, we analyzed the half-lives of K/R mutants expressed in Cx-deficient MDCK cells in which protein translation was blocked pharmacologically. Furthermore, we generated all possible double and single mutants of lysines 264, 287, and 303 (termed K264/303R, K264/287R, K287/303R, K264R, K287R, and K303R) to determine if one or more lysine residue become ubiquitinated and how many lysine residues needed to be mutated to inhibit GJ internalization. 24 hours after transient transfection protein translation was blocked by treatment with cycloheximide. Cells were lysed 0, 1, 2, 3, 4, and 6 hours post cycloheximide addition and total wild type and mutant Cx43 protein levels were analyzed using western blots (**Figure 4**). The half-life for wild type Cx43 was determined to be approximately 2.3 hours, correlating with the known, short 1-5 hours half-life described previously for Cx43 (Beardslee et al., 1998; Falk et al., 2009; Fallon and Goodenough, 1981) (**Figure 4A, top panel**). Unlike wild type, the protein levels of Cx43 in 6K/R and 3K/R at 6 hours post treatment had only decreased to 75% of starting Cx43 levels (**Figure 4A, middle and bottom panels)**. (The observed 25% loss of Cx43 protein observed in these cells over the 6 hour period is most likely due to a loss of cells caused by the cytotoxic effect of treating cells with cycloheximide, blocking protein biosynthesis in bulk, and associated negative metabolic side effects.) Using a linear fit, the half-lives of these mutants were extrapolated as 15.7 and 13.1 hours, respectively, making the 6K/R and 3K/R mutant half-lives 4-5 times longer than that of wild type Cx43. 6K/R and 3K/R mutant half-lives were not significantly different from one another, again suggesting that lysines 258, 345, and 346 are not targets for K63-polyubiquitination, and apparently play no direct role in GJ internalization. Expression of double-mutants (K287/303R, K264/303R, and K264/287R) resulted in a similar significantly increased half-live of all three mutants (app. 11.4, 6.6, and 7.7 hours, respectively), suggesting that apparently at least two separate lysine residues become K63-polyubiquitinated (**Figure 4B**). Finally, expressing the single lysine residue mutants also resulted in extended protein half-lives for K264 and K303 mutants (4.5 hours for each), while the half-life of the K287 mutant corresponded to the protein half-life of wild type Cx43 (2.5 hours) (**Figure 4C**). As already shown in **Figure 3C**, relevant mutants (all except the K287R mutant) showed significantly increased Cx43 protein levels compared to wild type Cx43 (**compare Figure 3C with Figure 4**). Taken together these results identified two juxtaposed lysine residues in the Cx43 C-terminal domain (K264 and K303) as target sites for K63-polyubiquitination and link K63-polyubiquitination to GJ internalization.

**Figure 4:**
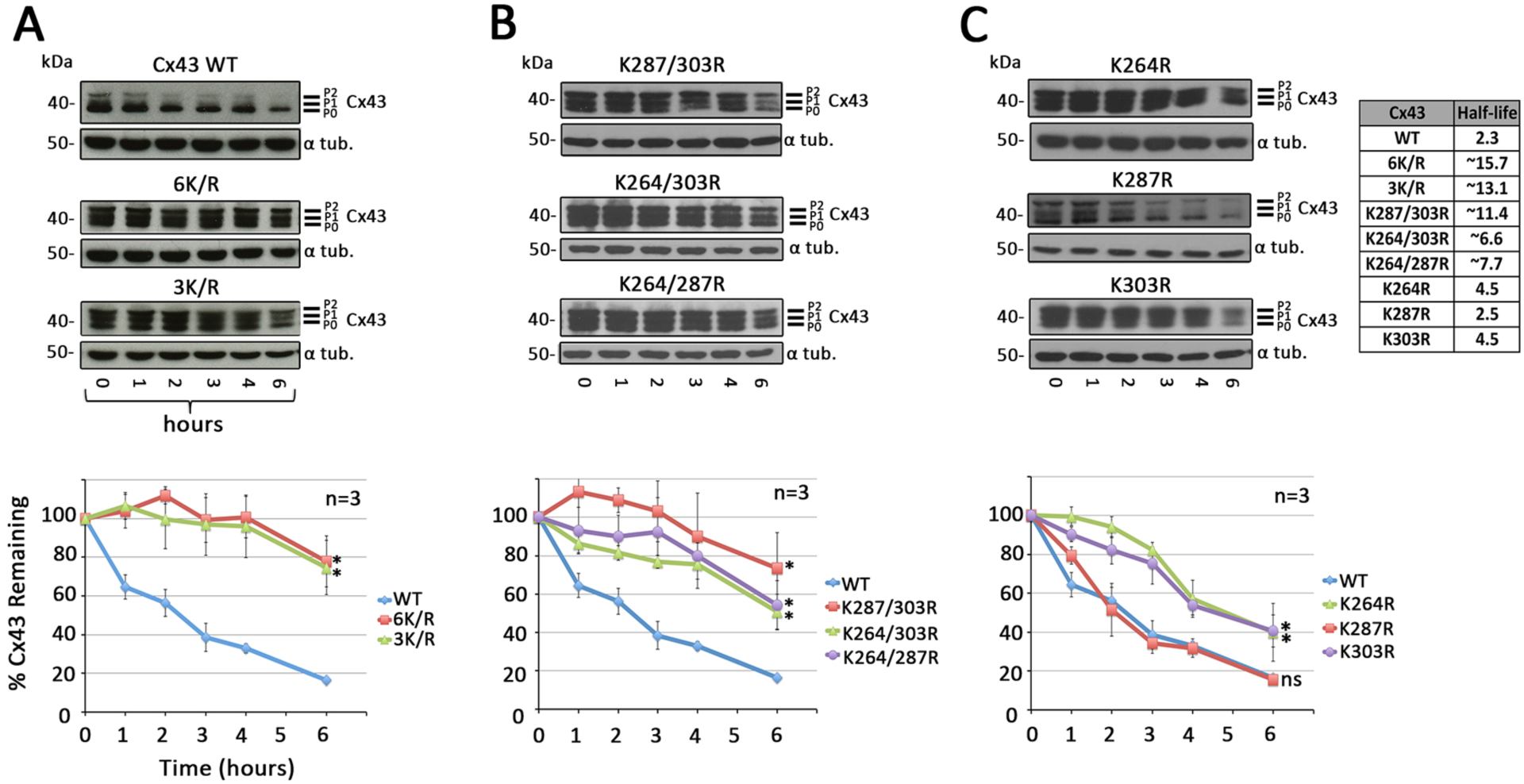
Cx43 K/R mutants with critical lysine residues mutated exhibit significantly extended protein half-lives. MDCK cells transfected with Cx43 wild type or K/R mutant constructs were treated with cycloheximide for indicated times, lysed and analyzed by western blot. **(A)** Quantification of Cx43 protein levels revealed a half-life of 2.3 hours for wild type Cx43, with app. 20% of the starting Cx43 protein remaining after 6 hours of treatment. Mutants 6K/R and 3K/R exhibited significantly increased half-lives extrapolated to 15.7 and 13.1 hours, respectively. **(B)** All double K/R mutants also exhibited significantly extended half-lives (see table in C). **(C)** Mutating lysine 264 and 303, together or independent of one another, also resulted in increased half-lives, whereas the half-live of the K287R mutant was unaffected and comparably short to wild type (2.5 hours). α-tubulin was probed as a loading control. Extrapolated half-lives were calculated using a linear fit curve.

### Immunoprecipitation of Cx43 mutants affirm K63-polyubiquitination on lysines 264 and 303

To further investigate whether lysines 264 and 303 are indeed the residues that become K63-polyubiquitinated in Cx43, we again expressed Cx43 wild type and relevant double and single K/R mutants (K264/303R, K264R, K287R, and K303R) in MDCK cells. Cells again were fractionated into soluble (Cx, connexon) and pellet fractions (GJ, AGJ) using TX-100 extraction (Musil and Goodenough, 1991). Then, wild type and mutant Cx43 polypeptides were immunoprecipitated from the pellet and probed for the presence of K63-polyUb modification with K63-polyUb-specific antibodies (**Figure 5**). Resulting Western blots showed that wild type, as well as the Cx43 K287R mutant could become K63-polyubiquitinated (**Figure 5A**) as indicated by the higher molecular weight band pattern that again included the weak, yet noticeable 70 and 150 kDa bands (marked with asterisks) in addition to a higher molecular weight smear (marked with bracket as detected previously under comparable conditions (compare Figure 3C, left panel with Figure 5A). In contrast, no higher molecular band patterns were detected in K264 and K303 double as well as single mutants, suggesting that these mutants did not become ubiquitinated; and that mutating one lysine may be sufficient to also prevent ubiquitination of the remaining lysine. Quantitative analysis of higher molecular weight Cx43 is shown in **Figure 5B**. Control lanes consisting of lysate plus Protein A-Sepharose beads, beads alone, or beads plus Cx43 antibody, as expected did not precipitate any K63-polyubiquitinated proteins. Amounts of total Cx43 and of K63-polyUb proteins (with α-tubulin as a loading control) present in the cell lysates were analyzed in control (**Figure 5C**). Taken together, these results further suggest lysines 264 and 303 as target lysines for K63-polyubiquitination of Cx43 in GJs. Our findings correlate with mass spectrometry evidence suggesting ubiquitination of Cx43 on lysines 9 and 303 (Wagner et al., 2011).

**Figure 5:**
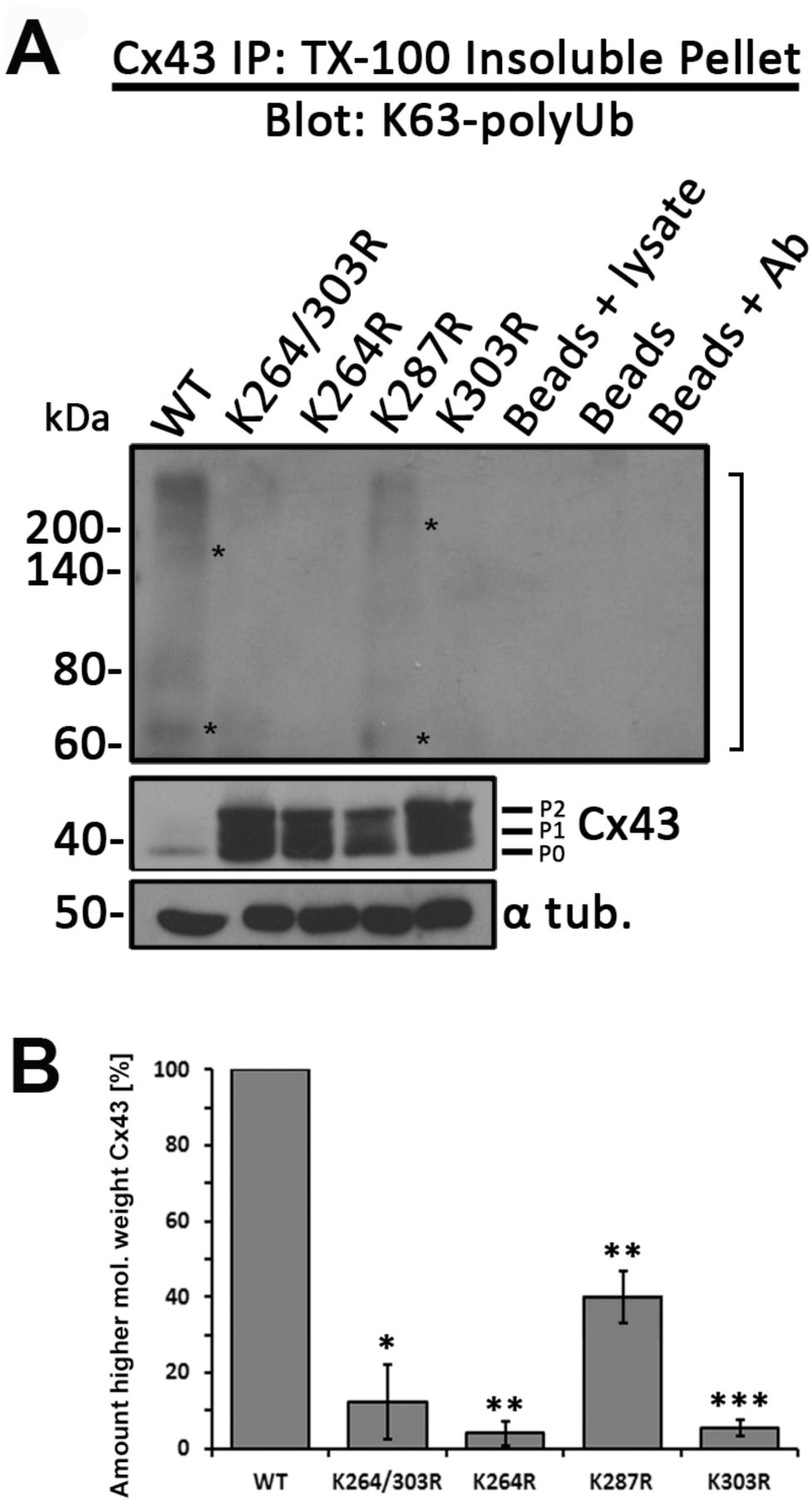
Cx43 K264R and K303R single and double mutants no longer become K63-polyubiquitinated. Cx43 was immunoprecipitated from MDCK cells transiently transfected with wild type, K264R, K287R, K303R and K264/303R mutants. Cells were lysed in 1% TX-100 containing buffer and separated into soluble and insoluble fractions. Cx43 was precipitated from insoluble fractions (GJs, AGJs) and probed for K63-polyUb by western blot as in Figure 2C. **(A)** Immunoprecipitation (and quantification in **B**) again revealed a higher-molecular weight smear (bracket) that included bands migrating at app. 70 and 150 kDa (asterisks) typical of ubiquitination in Cx43 wild type (compare Figure 2C, left panel) and in K287R mutant when probed with the K63-polyUb-specific antibodies. None of the other mutants revealed a similar higher molecular weight band pattern further suggesting that K264 and K303, but not K287 in Cx43 can become K63-polyubiquitinated. **(C)** Input cell lysates were probed in parallel for total K63-polyubiquitinated proteins (as in Figure 2B, right panel), Cx43, and α-tubulin as loading controls.

### Preventing ubiquitination on lysines 264 and 303 leads to hyper-phosphorylation of serines 368, 279/282, and 255

As described above, K63-polyubiquitination-deficient Cx43 polypeptides and GJs were found to accumulate in the plasma membrane. Intriguingly, these Cx43 mutants accumulated as hyper-phosphorylated variants (compare the amount of Cx43 P0 to P1 and P2 forms in Figures 3C and 4), suggesting that phosphorylation at specific sites in Cx43 may precede and trigger Cx43 ubiquitination. To further investigate which phosphorylation events preceded ubiquitination, we analyzed and quantified phosphorylation levels of the Cx43 lysine mutants on serines 368, 279/282, 255, and 262; phosphorylation events well known to down-regulate GJ mediated cell-to-cell communication (reviewed in Thevenin et al., 2013). MDCK cells were lysed 24 hours post transfection with wild type or relevant K/R mutant constructs, and analyzed by western blots using Cx43 phospho-specific antibodies (**Figure 6**). Amounts of phosphorylated serine 368 (pS368), 279/282 (pS279/pS282), and 255 (pS255) Cx43 polypeptides were significantly increased in all double and single K/R mutants with the exception of the non-ubiquitinated K287R mutant. Increased phosphorylation of serine 262 (pS262) was also observed, however only in a few mutants (some mutants containing 287, and/or 303 K/R exchanges), suggesting that mutating lysine 287 may leads to side effects that impair GJ internalization (discussed below). Taken together, these results indicate that in all Cx43 mutants that harbor lysine to arginine exchanges on K264 and/or K303, serines 368, 279/282, and 255 become hyper-phosphorylated; suggesting that phosphorylation on one, or a combination of these serine residues may trigger K63-polyubiquitination to induce GJ internalization.

**Figure 6:**
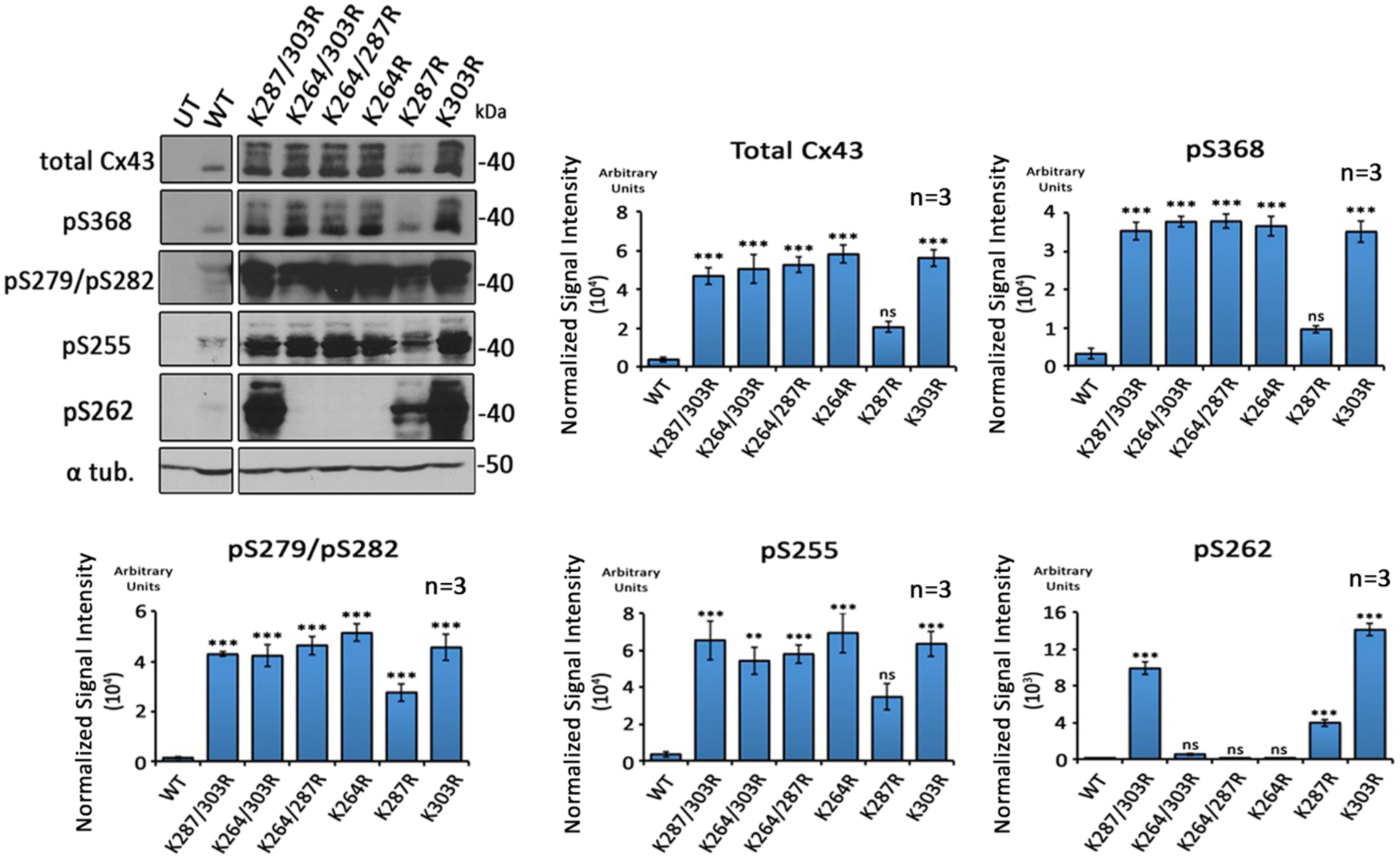
K63-polyUb-deficient Cx43 K/R mutants become hyper-phosphorylated on S368, S279/S282, and S255. Wild type, single and double K/R mutants expressed in MDCK cells were analyzed by western blot and probed using antibodies directed against total Cx43, and Cx43-phosphospecific antibodies directed against phosphorylated S368 (pS368), S279/S282 (pS279/pS282), S255 (pS255), and S262 (pS262) (specific Cx43 phosphorylation events known to down-regulate GJIC). α-tubulin was analyzed as loading control. Densitometric analyses of Cx43 proteins were normalized to α-tubulin. Dramatic increases in pS368, pS279/pS282, and pS255 were detected whenever K264 and/or K303 were mutated. Accumulation of Cx43 phosphorylated on serine 262 (pS262) was only detected in a few mutants (mutants containing K287R and/or K303R) and was attributed to potentially other, or unspecific mutagenesis-related side effects. Samples were analyzed on the same blots to allow direct comparison. Lanes of blots are shown separated when extraneous lanes were removed.

## DISCUSSION

Ubiquitination is a post-translational modification that plays an important role in regulating protein degradation. Particularly, the additions of K48- and K63-linked polyubiquitin chains are well-known signals for targeting proteins and protein-complexes to either proteasomal or endo-/phagolysosomal degradation (Komander and Rape, 2012). Here, we report for the first time the identification of two juxtaposed lysine residues in the C-terminus of Cx43, lysines 264 and 303, which apparently can become K63-polyubiquitinated in GJs. Cx43 lysine 264 and 303 to arginine exchange mutant proteins exhibited significantly longer protein half-lives and assembled significantly larger GJ plaques, suggesting that K63-polyubiquitination of Cx43 is required for successful GJ internalization. Our additional discovery that Cx43 ubiquitin-deficient lysine mutants accumulated as hyper-phosphorylated proteins in the plasma membrane further suggests that Cx43 K63-polyUb-mediated GJ internalization may be triggered by phosphorylation. Use of phospho-specific Cx43 antibodies revealed that hyper-phosphorylation occurred on serines 368, 279/282, and 255, well characterized PKC and MAPK targets known to down-regulate GJ-mediated intercellular communication (GJIC) ((Fong et al., 2014; Kanemitsu et al., 1998; Lampe et al., 1998; Lampe et al., 2000; Leykauf et al., 2003; Nimlamool et al., 2015; Petrich et al., 2002; Polontchouk et al., 2002; Saez et al., 1997; Sirnes et al., 2009; Solan and Lampe, 2007; Solan and Lampe, 2008) and GJ turnover (Fong et al., 2014; Nimlamool et al., 2015; (Thevenin et al., 2017).

In our experiments, K63-polyubiquitinated Cx43 migrated as a smear of higher molecular weight forms (marked with brackets in **Figures 2, 5**) including several distinct bands, e.g. 70 and ~150 kDa (marked with asterisks in **Figures 2, 5**) on SDS-PAGE gels similar in appearance as described by Ribeiro-Rodrigues and co-workers (Ribeiro-Rodrigues et al., 2014). Although ubiquitinated proteins are known to not necessarily migrate on SDS-PAGE gels according to their molecular weight (Hospenthal et al., 2015), polyubiquitin chains in general are believed to consist of at least 4 Ubs (Ravid and Hochstrasser, 2008). Four Ubs plus the molecular weight of Cx43 add up to 77 kDa, which is relatively close to the migration pattern of the in general prominently detected 70 kDa band, thus conforming to established criteria.

Ubiquitination of Cx43 in order to facilitate its degradation has been known for quite some time (Girao et al., 2009; Laing and Beyer, 1995; Laing et al., 1997; Leithe and Rivedal, 2004). Ubiquitination of Cx43 was discovered as a signal for proteasomal degradation (Laing and Beyer, 1995), thus likely regulating the degradation of mis-folded or aberrantly oligomerized Cx43 polypeptides shortly after their biosynthesis. The later discovery that ubiquitination may also play a role in the lysosomal degradation of Cx43 (Laing et al., 1997) suggested that ubiquitination may also play a role in the degradation of oligomerized Cx43 complexes (connexons, GJ channels, and GJ plaques). However, the type of ubiquitination, the lysine residues that become ubiquitinated, and the signals that may induce Cx43 ubiquitination in GJs to aid in their internalization and degradation remained illusive. It is now well known that the type of Ub linkage conveys Ub signal specificity (Ikeda et al., 2010; Komander and Rape, 2012; MacGurn et al., 2012). Specifically, exposure of a hydrophobic patch on Ub and flexibility of the Ub C-terminus rely on the type of polyUb chain conformation (e.g. K48-linked versus K63-linked polyUb chains) and thus determine Ub signal specificity (Ye et al., 2012). Different E3 ligases add ubiquitins in a chain specific manner. Nedd4-1, Wwp1, and Smurf2 (all HECT ligases) as well as Trim21 (a RING ligase) have been found to interact with Cx43 (Basheer et al., 2015; Chen et al., 2012; Fykerud et al., 2012; Leykauf et al., 2006). Of these, Nedd4-1 is known to preferentially interact with K63-polyubiquitinated proteins (Kim and Huibregtse, 2009) and to interact with ^283^PPGY^286^ in Cx43 via its WW2 domain (Leykauf et al., 2006) (see **Figure 7A**). Indeed, phosphorylation of S279/S282 *in vitro* was recently found to increase the binding affinity of this E3 ligase for its Cx43 binding domain (Spagnol et al., 2016), further supporting the concept that Nedd4-1 K63-polyubiquitinates Cx43 in GJs to mediate their internalization and degradation.

**Figure 7:**
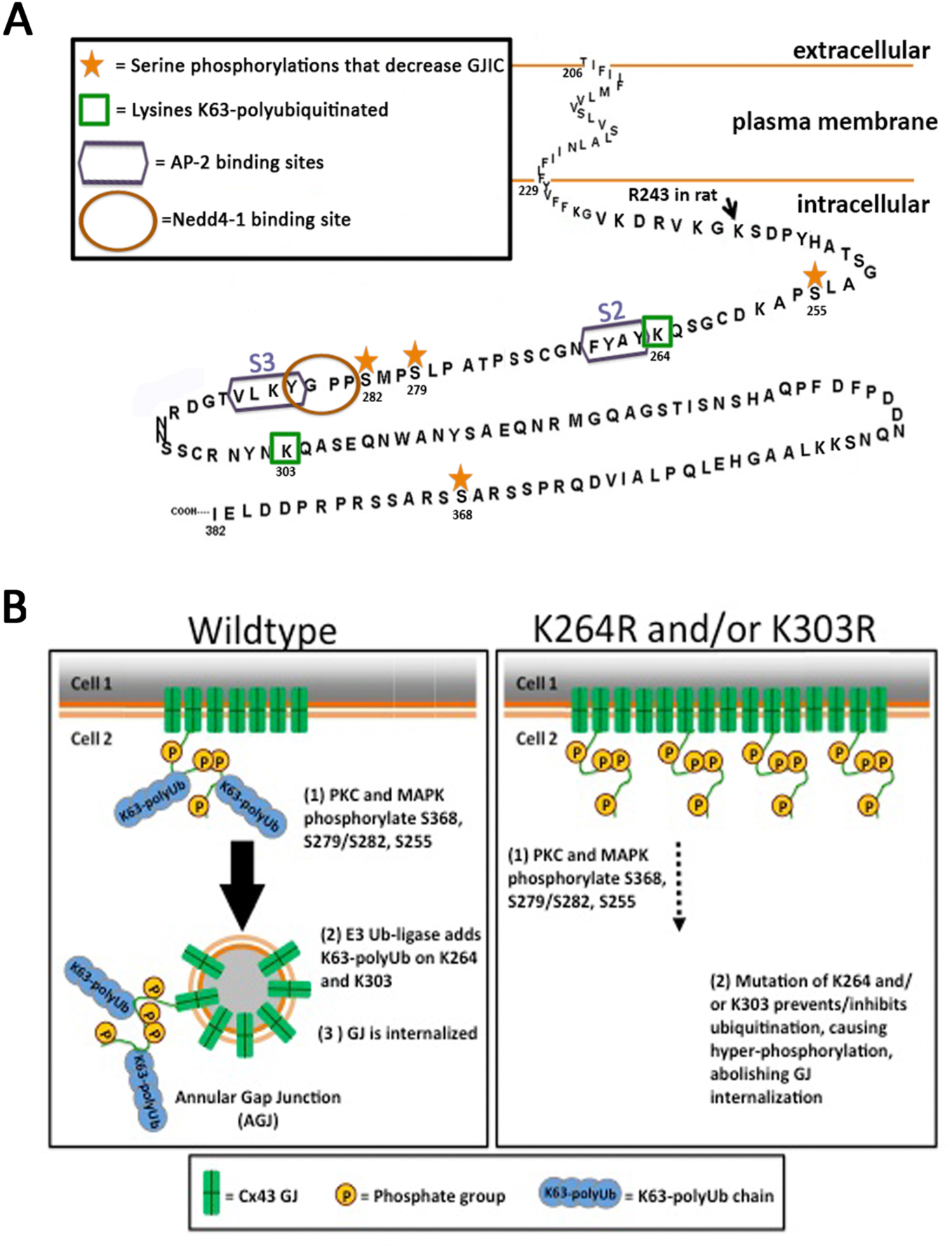
Schematic of K63-polyUb-mediated internalization of Cx43 GJs. **(A)** C-terminal tail of Cx43 showing the relevant Nedd4-1 E3-Ub ligase binding site (brown circle), the AP-2/clathrin binding sites (purple boxes), the K63-polyubiquitinated lysine residues (green boxes), and the serine residues that are phosphorylated preceding to Cx43 K63-polyubiquitination (golden asterisks). Note that ubiquitination occurs juxtaposed to the left and right of the Nedd4-1 and AP-2/clathrin binding sites. **(B)** Phosphorylation of Cx43 by PKC on S368, and MAPK on S279/S282, S255 and possibly on additional residues and kinases leads to Nedd4-1 recognition, K63-polyubiquitination, and AP-2/clathrin binding; resulting in GJ internalization. Mutating lysines 264 or 303 separately or together inhibits ubiquitination, causes Cx43 hyperphosphorylation on distinct residues and abolishes GJ internalization. Coupled S365 de-phosphorylation/S368 phosphorylation (termed gate-keeper event) may be critical, as it has been associated with a significant conformational re-arrangement of the Cx43 C-terminal domain that affects the upstream located, AP-2/clathrin binding region (Solan et al., 2007), and thus may allow MAPK, Nedd4-1 and AP-2/clathrin to access their respective Cx43-CT binding sites that otherwise may sterically be inaccessible in functional Cx43 GJ channels.

Cx43 domains exposed to the cytoplasm when oligomerized into GJ channels (intracellular/I-loop and C-terminal [CT] domain) contain a large number of lysine residues (11 in the I-loop, 9 in the CT) that are all potential targets for ubiquitination (labeled in red in **Figure 3A**). However, as the majority of regulatory phosphorylation events occur in the Cx43-CT, we hypothesized that ubiquitination - if occurring in GJs – would also occur in the Cx43-CT. In addition, the C-terminal region juxtaposed to the fourth transmembrane domain is known to harbor a microtubule binding site (Giepmans et al., 2001) and when mutated will likely interfere with Cx trafficking (Wayakanon et al., 2012). We thus left the lysines located within this region (K234, K237, and K241) untouched. Both mutants, with the 6 remaining C-terminal lysines (6K/R; K258, K264, K287, K303, K345, K346), and the mutant with the 3 central lysines mutated (3K/R; K264, K287, K303) behaved indistinguishable from each other, resulting in significantly increased mutant protein half-lives, the accumulation of hyper-phosphorylated Cx43 polypeptides, and larger GJ plaques in the plasma membrane (**Figures 3, 4**); suggesting that one or more of these 3 lysine residues become K63-polyubiquitinated. Generating and examining all possible double and single mutants then indicated that two of the three lysine residues, K264 and K303, can become K63-polyubiquitinated. Our finding that the protein half-lives of K264 and K303 single mutants were significantly longer compared to wild type, although not as long as half-lives of the double lysine mutants (4.5 hours and 6.6 −11.4 hours compared to 2.3 hours for wild type, **Figure 4C, Table**) argues for a cumulative effect. However, we did not detect K63-polyubiquitination of K264 and K303 single mutants (**Figure 5**), suggesting that mutating one lysine residue may already inhibit K63-polyubiquitination of Cx43 altogether. Of course, mutating lysines into arginines may result in conformational alterations of the Cx43-CT and this indirectly may inhibit GJ internalization, however the fact that exactly two out of six mutated lysines inhibited GJ internalization, that K63-polyubiquitination was detected on both locations (K264 and K303), and that the K287R mutant exhibited a half-life similar to wild type and was not hyper-phosphorylated strongly argues against this possibility.

Phosphorylation of Cx43 is well established and has been shown to be a regulatory mechanism for Cx43 trafficking, GJ assembly, gating, plaque internalization, and degradation (reviewed in Falk et al., 2016; Solan and Lampe, 2014; Thevenin et al., 2013). Cx43 phosphorylation by Akt (protein kinase B), PKA (protein kinase A), and CK1 (casein kinase 1) is known to up-regulate GJ intercellular communication (GJIC), whereas phosphorylation by PKC (protein kinase C), CDC2 (cell division cycle protein 2), MAPKs, and Src down-regulate GJIC by closing GJ channels (Fong et al., 2014; Kanemitsu et al., 1998; Lampe et al., 1998; Lampe et al., 2000; Leykauf et al., 2003; Nimlamool et al., 2015; Petrich et al., 2002; Polontchouk et al., 2002; Saez et al., 1997; Sirnes et al., 2009; Solan and Lampe, 2007; Solan and Lampe, 2008; van Zeijl et al., 2007). More recent findings from our lab also provided evidence for a link between Cx43 phosphorylation and GJ internalization. Specifically, phosphorylation of serines 368, 279/282, 255 (and in some cases 262) by PKC and MAPKs in response to epidermal-(EGF) and vascular endothelial growth factor (VEGF) stimulation has been linked to acute GJ internalization in mouse embryonic stem cells and pPAECs (Fong et al., 2014; Nimlamool et al., 2015); and on serines 365, 368 and 373 to regulate Cx43/ZO-1 binding and release, an important early step in GJ turnover (Thevenin et al., 2017). Additionally, S279/S282 phosphorylation-deficient Cx43 mutants were shown to stabilize GJ plaques in human pancreatic tumor cells (Johnson et al., 2013), and upon EGF stimulation in HeLa cells (Schmitt et al., 2014). Recently, Solan and Lampe (Solan and Lampe, 2015) and we (Falk et al., 2016) suggested a kinase program consisting of PKC, MAPKs, and Src that hierarchically phosphorylate Cx43 on serines 368, 279/282 (and potentially 255 and 262), and tyrosine 247, respectively to spatiotemporally regulate GJ internalization. Of these events, phosphorylation of S368 is of particular interest. It has been found to preferentially localize to the center of GJ plaques (Cone et al., 2014), the region where GJ channel internalization occurs (Falk et al., 2009; Gaietta et al., 2002; Lauf et al., 2002). Furthermore, S368 phosphorylation is known to be dependent on S365 dephosphorylation (termed “gate keeper”), an event known to trigger a large conformational change of the Cx43-CT that affects the upstream region that harbors the above described ubiquitination sites (Solan et al., 2007). However a link between phosphorylation and ubiquitination has not been demonstrated.

Ub-mediated protein degradation is often regulated via protein phosphorylation (termed phosphodegron). Phosphorylation on one or more residues induces substrate changes (conformational or other) allowing the subsequent ubiquitination of target lysines that then triggers the degradation of the phosphorylated/ubiquitinated protein (Nguyen et al., 2013; Ravid and Hochstrasser, 2008). As our data suggests that phosphorylation of S368, S279/S282, and S255 occurs prior to ubiquitination, and preventing ubiquitination on relevant lysine residues results in the build-up of hyper-phosphorylated mutant Cx43 protein and GJs in the plasma membrane, it is possible that similarly, phosphorylation of Cx43 induces K63-polyubiquitination that then triggers GJ internalization and degradation. We do not know yet which, if any of the three phosphorylation events are required and in which order. However, as S365 de-phosphorylation/S368 phosphorylation is linked to conformational Cx43-CT rearrangements (Solan et al., 2007), and S279/S282 phosphorylation is linked to enhancing Nedd4-1 binding (Spagnol et al., 2016), phosphorylation on at least these two sites and in this order is likely to precede Cx43 K63-polyubiquitination.

How might K63-polyubiquitination of Cx43 induce GJ internalization and degradation? It should be noted that at any given time we detected only a small portion of plasma membrane localized, TX-100 resistant Cx43 to be K63-polyubiquitinated. This indicates, that ubiquitination is a transient event, and only occurs on a small number of Cx polypeptides in GJ plaques. We know that cells can turn over GJs in two ways, (i) by continuously internalizing small GJ plaque portions preferentially from central plaque areas (Falk et al., 2009), and (ii) by internalizing entire plaques or large portions of plaques (Piehl et al., 2007). It is likely that in both instances, only critical connexin polypeptides need to become ubiquitinated (located in the periphery of the invaginating plaque area) to initiate internalization, while secondary MAPK-mediated phosphorylation events on S279/282, S255 (and eventually S262), that all are located in the vicinity of the two AP-2 binding sites that we characterized in the Cx43 C-terminal domain (Y^265^AYF and Y^286^KLV) (Fong et al., 2013), recruit clathrin and internalize GJs and GJ plaque portions (reviewed in Falk et al., 2016) (**Figure 7B**). In addition, unmodified Cx43 migrating at its predicted molecular weight of 43 kDa precipitated together with ubiquitinated Cx43 when K63-polyubiquitinated proteins were precipitated (**Figure 2C**, labeled with Cx43), suggesting that only a portion of Cx subunits in a connexon become ubiquitinated.

In addition, under certain conditions clathrin may be recruited by another group of adaptor proteins, termed clathrin associated sorting proteins, or CLASPs that specifically bind via a Ub-interacting motif (UIM) to a polypeptide sequence that is exposed in K63-polyubiquitin chains (Traub and Bonifacino, 2013). One such alternative clathrin adaptor, called Eps15, was proposed to bind to Cx43 and facilitate GJ internalization (Catarino et al., 2011; Girao et al., 2009). Thus, K63-polyubiquitination may allow Eps15 to bind to Cx43 and internalize GJs alternative to AP-2 (Gumpert et al., 2008; Piehl et al., 2007) (Falk et al., 2016; Falk et al., 2014; Gumpert et al., 2008; Piehl et al., 2007). Previously, we and others also have demonstrated that Cx43 in GJs and internalized GJs interacts with a protein termed p62/SQSTM1, which sequesters AGJs to autophagosomal degradation (Bejarano et al., 2012; Fong et al., 2012; Lichtenstein et al., 2010). p62/SQSTM1 interacts specifically with K63-polyubiquitinated substrates via its UBA (ubiquitin associated) domain (Seibenhener et al., 2004) to sequester targets for autophagosomal degradation (Bjorkoy et al., 2005; Pankiv et al., 2007). Depleting cells of autophagy-associated proteins such as LC3, Beclin1, Atg5/Atg12, and p62/SQSTM1 by RNAi significantly impaired GJ degradation (Bejarano et al., 2012; Fong et al., 2012; Lichtenstein et al., 2010). Thus, it is also likely that Cx43 K63-polyubiquitination sequesters internalized GJs for autophagosomal degradation.

Taken together, our current and previous findings suggest that a serious of post-translational modifications including phosphorylations/de-phosphorylations, ZO-1 binding and release, and K63-polyubiquitination transition functional into non-functional GJ channels that then are endocytosed (Fong et al., 2014; Nimlamool et al., 2015; Thevenin et al., 2017; reviewed in Falk et al., 2016). Blocking K63-polyubiquitination apparently prevents efficient GJ internalization and leads to the accumulation of hyper-phosphorylated Cx43 and GJs in the plasma membrane (schemed in **Figure 7B**). As E3-Ub specific small molecule inhibitors are available, clinical administration may prevent the acute loss of GJs from intercalated discs as is typical for a number of severe heart diseases (Fontes et al., 2012).

## MATERIALS AND METHODS

### 1 cDNA constructs and mutagenesis

Full-length wild type rat Cx43 cDNA was cloned into pEGFP-N1 vector (Clontech) as described (Falk, 2000). To generate full length untagged wild type Cx43, an authentic TAA stop codon was re-introduced (Fong et al., 2013). Untagged rat Cx43 mutant 3K/R and 9K/R cloned into pcDNA3.1 were generously provided by Vivian Su and Alan Lau (Natural Products and Cancer Biology Program, Cancer Research Center of Hawaii, Honolulu, HI 96813) (Su et al., 2010). Untagged 6K/R Cx43 mutant was generated by restoring K234, K237, and K241 from 9K/R and adding a BglII restriction site for confirmation of mutagenesis. Double and single K/R mutants were generated by restoring one or two of the remaining lysines 264, 287, and 303 from the Cx43 mutant 3K/R using Quick Change Mutagenesis (Stratagene, Santa Clara, CA). Forward recovery mutagenesis primers were as follows (arginine recovered codons are underlined, BglII site in bold): **R234_237_241K recovery**, 5’-G CTC TTC TAC GTC TTC TTC <u>AAA</u> GGC GTT <u>AAG</u> GAT CGC GTG <u>AAG</u> GGA **AGA TCT** GAT CC-3’; **R264K recovery**, 5’-CCA TCA <u>AAA</u> GAC TGC GGA TCT CCA AAA TAC GCC TAC TTC AAT GGC-3’; **R287K recovery**, 5’-CCT ATG TCT CCT CCT GGG TAC <u>AAG</u> CTG GTT ACT GGT GAC AGA AAC AAT TCC-3’; and **R303K recovery**, 5’-CC TCG TGC CGC AAT TAC AAC <u>AAG</u> CAA GCT AGC GAG CAA AAC TGG-3’. PCR mutagenesis reactions were generated using proofreading *Pfu* Ultra II polymerase (Cat. No. 600670-51; Stratagene). Digestion of template cDNA with *Dpn1* restriction endonuclease (Cat. No. R0176-S; New England Biolabs, Ipswich, MA) was followed by transformation into chemically competent DH5α *E. coli* cells (Cat. No. 18258-012; Invitrogen). Sequencing was performed on all cDNA constructs to confirm the presence of intended mutations and exclude the presence of additional unwanted mutations.

### 2 Cell culture and transient transfections

HeLa (gap junction deficient; Cat. No. CCL-2; American Type Culture Collection, Manassas, VA), Madine-Darby Canine Kidney (MDCK) (gap junction deficient; Cat. No. NBL-2; American Type Culture Collection), and primary porcine Pulmonary Artery Endothelial Cells (pPAECs) (expressing Cx43 and other Cxs, gap junction proficient; Cat. No. P302; Cell Applications, San Diego, CA) were maintained in low glucose Dulbecco’s Modified Eagle Medium (DMEM) (Cat. No. SH30021.01; HyClone, Logan, UT) supplemented with 50 I.U/ml penicillin and 50μg/ml streptomycin (Cat. No.30-001-Cl; Corning, Manassas, VA), 2mM Lglutamine (Cat. No. 25-005-C1; Mediatech, Manassas, VA), and 10% Fetal Bovine Serum (Cat. No. S11150; Atlanta Biologicals, Flowery Branch, GA) at 37°C, 5% CO_2_, and 100% humidity. Cells were washed with 1x PBS and treated with 0.25% trypsin (Cat. No. 25-053-Cl; Corning) for passaging. 24-48 hours after passaging, 60-80% confluent HeLa and MDCK cells were transiently transfected with wild type and mutant constructs using Superfect (Cat. No. 301307; Qiagen, Hilden, Germany) or Lipofectamine2000 (Cat. No. 11668019; Invitrogen, Carlsbad, CA) reagents, respectively, according to manufacturer’s recommendations.

### 3 Immunofluorescence staining and image analyses

pPAECs and HeLa cells were grown on pretreated poly-L-Lysine (Cat. No. P8920; Sigma, St. Louis, MO) coated coverslips in low glucose DMEM at 37°C, 5% CO_2_, and 100% humidity. Cells were fixed in 3.7% formaldehyde/PBS and permeabilized with 0.2% Triton X-100 (TX-100) (Cat. No. 3929-2; VWR, Radnor, PA) in PBS. Cells were blocked in 10% FBS/PBS for 30 minutes at room temperature (RT) and incubated with primary rabbit polyclonal anti-peptide Cx43 antibodies (Cat. No. 3512; Cell Signaling, Danvers, MA) or rabbit monoclonal anti-Cx43 antibodies (clone 4E6.2) (Cat. No. MAB3067, Millipore, Billerica, MA) diluted 1:500 in 10% FBS/PBS at 4°C overnight. Additionally, pPAECs were incubated with primary mouse monoclonal anti-monoUb and polyUb (FK2) (Cat. No. PW8810; Enzo, Farmingdale, NY), polyUb (FK1) (Cat. No. PW8805; Enzo), K63-polyUb (clone HWA4C4) (Cat. No. PW0600; Enzo), or primary monoclonal K48-polyUb (clone Apu2) (Cat. No. 05-1307; Millipore) antibodies diluted 1:200 in 10% FBS/PBS at 4°C overnight. Cells were incubated in secondary antibodies (goat anti-rabbit Alexa Fluor488, goat anti-mouse Alexa Fluor568, goat anti-rabbit Alexa Fluor568, and goat anti-mouse Alexa488; Cat. No. A11008, A11031, A11011, and A11001, respectively; Molecular Probes/Invitrogen, Eugene, OR) diluted 1:200 or 1:500 in 10% FBS/PBS, respectively, for 1 hour at RT. Cells were also stained with 1 μg/ml 4′,6-diamidino-2-phenylindole (DAPI) (Cat. No. D1306; Molecular Probes). Coverslips were rinsed in distilled water and mounted using Fluoromount-G (Cat. No. 0100-01; Southern Biotechnology, Birmingham, AL). pPAEC and HeLa cells were imaged using a Nikon Eclipse TE2000E wide-field inverted fluorescence microscope equipped with 40x NA 1.3 Plan Fluor and 60x NA 1.4 Plan-Apochromat oil-immersion objectives (Nikon Instruments, Melville, NY) and a forced-air cooled Photonics CoolSnap^®^ HQ CCD camera (Roper Scientific, Duluth, GA). Images were acquired using MetaVue software version 6.1r5 (Molecular Devices, Sunnyvale, CA). To quantify GJ plaque size, plaques were outlined using the freeform tool in ImageJ (National Institutes of Health, Bethesda, MD) and total pixel number/outlined area was quantified for 73, 98 and 83 cell pairs, expressing Cx43 wild type, 6K/R, and 3K/R, respectively. Similarly, to quantify colocalization of ubiquitin-specific antibodies with Cx43 antibodies, GJ plaques of entire lateral membranes were outlined (at least 6/antibody/image), and pixel intensity determined. Next, the signal received using the colocalization function in ImageJ was determined and compared to total Cx43 fluorescence to determine percent colocalization. Different threshold settings were chosen and quantified, however all chosen ones always resulted in zero percent colocalization between Cx43 and Apu2 (K48-polyUb specific) antibodies.

### 4 Triton X-100 solubility assays

To separate intracellular Cx proteins and connexons from plasma membrane GJs and likely also internalized AGJs, TX-100 solubility assays were performed based on the method described by Musil and Goodenough (Musil and Goodenough, 1991). MDCK cells transiently transfected with wild type or Cx43 K/R constructs or pPAECs were used. 24 hours post transfection (MDCKs), or 48 hours after seeding (pPAECs), cells were washed in ice cold PBS. Prior to lysis, pPAECs were treated with 20 μM of the pan-DUB inhibitor, PR-619, (Cat. No. 662141; EMD Millipore) for 1.5 hours (Figure 2B). Cells were lysed on ice for 15 min in 400 μL pre-chilled lysis buffer containing 50 mM Tris-HCl, pH7.4 (Cat. No. BP 153-1; Fisher Bioreagents, Fair Lawn, NJ), 150 mM NaCl (Cat. No. S671-3; Fisher Chemicals, Fair Lawn, NJ), 1 mM EDTA (Cat. No. 161-0729; Biorad, Hercules, CA), 0.5% sodium deoxycholate (Cat. No. D6750; Sigma), 1% Triton X-100 (Cat. No. VW3929-2; VWR), and 1% Igepal (Cat. No.18896; Sigma). The buffer was supplemented with the following protease and phosphatase inhibitors: 1 mM protease inhibitor cocktail (Cat. No. P8340; Sigma), 1 mM β-glycerol phosphate (Cat No. 157241; MP Biomedicals, LLC, Solon, OH), 1 mM sodium orthovanadate (Cat. No. 450234; Sigma), 20 mM N-ethylmalemide (Cat. No. E1271; Sigma), and 10 mM 1,10-phenanthroline monohydrate (Cat. No. AC130130050; ACROS Organics, NJ). Lysates were centrifuged at 10,000xg for 5 min at RT to pre-clear cell lysates. 180 μL of each resulting supernatant was centrifuged at 100,000xg in a Beckman Coulter Airfuge^®^ ultracentrifuge for 10 minutes. For western blot analysis, 100,000xg supernatants (TX-100 soluble fractions) and pellets (TX-100 insoluble fractions) were re-suspended in SDS-PAGE sample buffer and boiled for 5 minutes.

### 5 SDS-PAGE and Western blot analyses

Immunoprecipitation, protein half-life, and phosphorylation analyses samples were loaded onto 10% SDS-PAGE mini-gels (Biorad). Proteins were transferred onto nitrocellulose membranes and blocked for 1 hour at RT in 5% non-fat dry milk/TBST or 5% bovine serum albumin (BSA) (Cat. No. A7906; Sigma)/TBST. Antibodies were diluted in 5% BSA/TBS as follows: rabbit anti-Cx43, rabbit anti-K63-polyUb (Cat. No. 05-1308; EMD Millipore), rabbit anti-Cx43 pS279/pS282, rabbit anti-Cx43 pS255, rabbit anti-Cx43 pS262 (Cat. No. sc-12900-R, sc-12899-R and sc-17219-R, respectively; Santa Cruz, Dallas, TX) and rabbit anti-Cx43 pS368 (Cat. No. 3511S; Cell Signaling) at 1:2000, and mouse anti-α-tubulin primary antibodies at 1:5000. Blots were incubated with primary antibodies at 4 °C overnight, then washed three times with TBST. Secondary HRP-conjugated goat anti-rabbit or goat anti-mouse antibodies (Cat. No. 81-6520 or 81-6120, respectively; Zymed, San Francisco, CA) were diluted 1:5000 and secondary HRP-conjugated mouse anti-rabbit light chain specific antibodies (Cat. No. 211032-171; Jackson ImmunoResearch Laboratories, Inc., West Grove, PA) were diluted 1:15,000. Blots were incubated in secondary antibodies for 1 hour at RT. Proteins were detected using Pierce^®^ ECL2 Western Blotting Substrate (Cat. No. 80196; Thermo Scientific, Rockford, IL) and Kodak BioMax^®^ Light Autoradiography Film (Cat. No. 1788207; Carestream Health, Rochester, NY). Cx43 protein amounts were quantified using ImageJ software and normalized to α-tubulin.

### 6 Cx43 protein half-life analyses

24 hours post transfection, MDCK cells were treated with 50 μg/ml cycloheximide (Cat. No. C655; Sigma) for 0, 1, 2, 3, 4, and 6 hours at 37 °C, 5% CO_2_ and 100% humidity. Cells were lysed in SDS sample buffer at each time point and boiled for 5 minutes. Samples were separated on 10% SDS-PAGE gels using western blot protocols. Blots were probed with rabbit anti-Cx43 antibody, then stripped with stripping buffer and stringently washed in TBS/0.1% Tween20 (Cat. No. BP337; Fisher Bioreagents) (TBST) before re-probing with mouse anti-α tubulin antibodies (Cat. No. 9026; Sigma). Cx43 protein intensities were quantified using ImageJ and normalized to the corresponding α-tubulin intensities.

### 7 Immunoprecipitations

100,000xg pellets (TX-100 insoluble fraction) from MDCK cells transiently transfected with Cx43 wild type or K/R mutants were resuspended in fresh lysis buffer without TX-100 and sonicated (six 1 second/pulses), with 5-minute intervals on ice in between pulses. Immunoprecipitation samples were incubated with rabbit anti-Cx43 or mouse anti-K63-polyUb antibodies adsorbed to protein A-Sepharose beads (Cat. No. 3391; Sigma) for 2 hours at RT while rotating. After incubation, beads were washed in lysis buffer and immunoprecipitated protein was eluted from beads with SDS sample buffer and boiled for 5 minutes.

### 8 Statistical analyses

One-way ANOVA analyses with Post-Hoc *Bonferroni* corrections were performed on data sets including GJ plaque size measurements (**Figure 3B**), Cx43 wild type and mutant protein levels (**Figure 3C**), and amounts of phosphorylated Cx43 in K/R mutants (**Figure 6**) in SPSS software. Unpaired two-tailed student’s t-tests were performed in Excel to analyze Cx43 protein half-lives (**Figure 4**) and amounts of higher molecular weight K63-polyUb immunoreactive Cx43 (**Figure 5C**). All data are presented as mean ±SEM. p-values of *p<0.05, **p<0.005, and ***p<0.0005 were considered statistically significant.

## ACKNOWLEDGEMENTS

We thank Vivian Su and Alan Lau (Natural Products and Cancer Biology Program, Cancer Research Center of Hawaii, Honolulu, HI 96813) for providing Cx43 3K/R and 9K/R constructs, Lynne Cassimeris (Department of Biological Sciences, Lehigh University, Bethlehem, PA 18015) for the use of specialized equipment (AirFuge), JoAnn Trejo and Neil Grimsey, and current and previous Falk lab members for constructive discussions. This work was supported by NIHs NIGMS grant GM55725 to MMF.

## COMPETING INTERESTS

The authors declare that they have no conflict of interest.

## AUTHOR CONTRIBUTIONS

RMK-A and MMF designed the research, analyzed the data, and wrote the manuscript. RMK-A, RAM, and CGF performed experiments and analyzed data.

